# Optimization of graph construction can significantly increase the power of structural brain network studies

**DOI:** 10.1101/553743

**Authors:** Eirini Messaritaki, Stavros I. Dimitriadis, Derek K. Jones

## Abstract

Structural brain networks derived from diffusion magnetic resonance imaging data have been used extensively to describe the human brain, and graph theory has allowed quantification of their network properties. Schemes used to construct the graphs that represent the structural brain networks differ in the metrics they use as edge weights and the algorithms they use to define the network topologies. In this work, twenty graph construction schemes were considered. The schemes use the number of streamlines, the fractional anisotropy, the mean diffusivity or other attributes of the tracts to define the edge weights, and either an absolute threshold or a data-driven algorithm to define the graph topology. The test-retest data of the Human Connectome Project were used to compare the reproducibility of the graphs and their various attributes (edges, topologies, graph theoretical metrics) derived through those schemes, for diffusion images acquired with three different diffusion weightings. The impact of the scheme on the statistical power of the study and on the number of participants required to detect a difference between populations or an effect of an intervention was also calculated.

The reproducibility of the graphs and their attributes depended heavily on the graph construction scheme. Graph reproducibility was higher for schemes that used thresholding to define the graph topology, while data-driven schemes performed better at topology reproducibility. Additionally, schemes that used thresholding resulted in better reproducibility for local graph theoretical metrics, while data-driven schemes performed better for global metrics. Crucially, the number of participants required to detect a difference between populations or an effect of an intervention could change by a large factor depending on the scheme used, affecting the power of studies to reveal the effects of interest.

## 1 Introduction

The view that the human brain is a network of cortical and subcortical areas connected via white matter tracts emerged several decades ago and has been very useful in characterizing brain structure and function in both healthy participants and diseased populations (Hagmann et al. (2008); Griffa et al. (2013); Caeyenberghs and Leemans (2014); Fischer et al. (2014); van den Heuvel and Fornito (2014); Yuan et al. (2014); Baker et al. (2015); Collin et al. (2016); Drakesmith et al. (2015); Aerts et al. (2016); Nelson et al. (2017); Vidaurre et al. (2018); Imms et al. (2019), to name just a few of the studies). Structural networks can be constructed using data collected non-invasively via magnetic resonance imaging (MRI) and diffusion MRI (dMRI). Graph theory allows quantification of the organizational properties of structural brain networks by using graphs to represent those networks (Bullmore and Sporns (2009) and references therein).

For inferences drawn from graph theoretical analyses of structural brain networks to be reliable, it is essential that the graphs representing the structural networks reflect the true structural organization of the brain. If that is the case, such graphs are bound to be reproducible, within experimental error, when generated from data collected at different times, in the absence of any true changes in the structural connectome. Conversely, this means that graphs that are not reproducible in the absence of age-related changes or intervention-induced plasticity are not reliable in representing the structural organization of the human brain. This is important because various graph construction schemes have been presented in the literature that use different techniques to perform tractography, define graph topology and assign edge-weights, all producing graphs that are quite different from each other, with different levels of robustness and reproducibility. Using schemes that result in highly reproducible graphs and graph-attributes means that any observed changes (in longitudinal studies) or differences between populations (in comparative studies) can be reliably attributed to maturation or to differences between populations respectively, rather than to random fluctuations resulting from experimental or analysis errors. Additionally, high reproducibility and low within-participant variability results in higher power for the studies and therefore a lower number of required participants, which in turn results in reduction of the cost and resources required for the study. Several papers have investigated different aspects of the the reproducibility of structural brain networks and their graph theoretical metrics (Dennis et al., 2012; Owen et al., 2013; Buchanan et al., 2014; Smith et al., 2015; Owen et al., 2013; Zhong et al., 2015; Dimitriadis et al., 2017b; Yuan et al., 2018), each using a specific scheme for constructing the graphs, predominantly using data acquired with low diffusion weightings (*b*-values of up to 1500 s/mm^2^). The work of Roine et al. (2018) also investigated a higher diffusion weighting of *b* = 3000 s/mm^2^.

In this work, we compared the reproducibility of structural brain networks generated with different graph-construction schemes for three different diffusion weightings. We investigated the reproducibility of the graphs themselves as well as their topologies, edge-weights and graph theoretical metrics. The article is organized as follows: Section 2 details the data, the software and algorithms used to perform the tractography, construct the graphs representing the structural networks, and perform the comparative analyses. Section 3 contains a comparison of the graphs derived using different graph-construction schemes and describes the results for the reproducibility of the graphs and their various attributes. Finally, it presents sample calculations for the number of participants required for different studies, for the different graph-construction schemes. Section 4 includes a discussion of the significance of our results and the ways they can impact graph theoretical studies of structural brain networks in the future. We conclude with Section 5. To the best of our knowledge, this is the first time that graph-construction schemes for structural brain networks have been compared in terms of their reproducibility across a wide range of diffusion weightings, in particular with an aim to inform decisions on the number of participants required in related studies.

## 2 Methods

All analyses were performed using MATLAB (MATLAB and Statistics Toolbox Release 2015a, The MathWorks, Inc., Massachusetts, Unites States), unless otherwise stated.

### 2.1 Data

We used the Human Connectome Project (HCP) (Essen et al., 2013; Sotiropoulos et al., 2013b; Glasser et al., 2013) test-retest MRI and diffusion-MRI data, in which participants were scanned twice. The diffusion-weighted images (DWIs) have resolution of (1.25 × 1.25 × 1.25) mm^3^, and were acquired for three different diffusion weightings (*b*-values: 1000 s/mm^2^, 2000 s/mm^2^ and 3000 s/mm^2^). We used the data from 37 participants for whom there were 90 gradient directions for each *b*-value. The HCP acquisition details are described in Sotiropoulos et al. (2013b); Feinberg et al. (2010); Moeller et al. (2010); Setsompop et al. (2012); Sotiropoulos et al. (2013a); Xu et al. (2012). We performed the analyses described below separately for the DWIs collected with each diffusion weighting.

### 2.2 Tractography

We performed whole-brain tractography using ExploreDTI-4.8.6 (Leemans et al., 2009). Constrained Spherical Deconvolution (CSD) (Tournier et al., 2004) was used to estimate the fiber orientation distribution function. In the tractography algortihm, the seed point resolution was (2 × 2 × 2 mm^3^), the step size was 1 mm, the angle threshold was 30°, and the fiber length range was 50 − 500 mm.

### 2.3 Graph Generation

We constructed graphs using twenty different graph-construction schemes. We normalized all graphs so that the maximum edge weight in each graph was equal to 1. We also set the elements of the diagonal of all graph matrices equal to 0, since they correspond to connections of a node with itself.

#### 2.3.1 Node definition

We used the Automated Anatomical Labeling (AAL) atlas (Tzourio-Mazoyer et al., 2002) to define the 90 cortical and subcortical areas of the cerebrum that correspond to the nodes of the structural networks. The white matter (WM) tracts linking those areas are the connections, or edges, of the networks. The network generation was performed in ExploreDTI-4.8.6 (Leemans et al., 2009). This process resulted in nine connectivity matrices (CMs) for the data for each diffusion weighting and each scan of each participant. Each CM had edges weighted by a different metric averaged along the corresponding WM tracts. Those metrics are: fractional anisotropy (FA), mean diffusivity (MD), radial diffusivity (RD), number of streamlines (NS), streamline density (SLD), percentage of streamlines (PD), tract volume (TV), tract length (TL) and Euclidean distance between the nodes (ED), and are listed in Table 1.

**Table 1:** Metrics used in connectivity matrices.

| **Metric** | **Abbreviation** |
| --- | --- |
| Fractional anisotropy | FA |
| Mean diffusivity | MD |
| Radial diffusivity | RD |
| Number of streamlines | NS |
| Percentage of streamlines | PS |
| Streamline density | SLD |
| Tract volume | TV |
| Tract length | TL |
| Euclidean distance between nodes | ED |

#### 2.3.2 Integrated graphs

We used the algorithm described by Dimitriadis et al. (2017b, c) to generate an integrated network for the data from each diffusion-weighting and each scan of each participant. Following Dimitriadis et al. (2017b), for each participant at each time point, a two-step process was followed:

1. We used the diffusion-distance (Hammond et al., 2013) between individual CMs to maximise information provided by each metric and create a linear-combination (or integrated) graph, and
2. we used an Orthogonal Minimal Spanning Tree (OMST) algorithm (Dimitriadis et al., 2017a, c) to selectively remove edges of the resulting graph, so as to maximize the difference (Global Efficiency - Cost), while maintaining the connectivity of the nodes. The benefit of the method lies in the fact that both the topology that results from selectively removing edges and the assignment of the edge weights are performed in a data-driven manner, and no arbitrary threshold needs to be imposed. This additionally ascertains that both strong and weak edges are treated equally.

Motivated by the fact that the nine metrics shown in Table 1 exhibit a number of covariances, we also considered integrated graphs formed from subsets of those metrics, to explore whether using fewer variables to define edge-weights has an impact on network reproducibility.

#### 2.3.3 Single-metric graphs

In addition to using the algorithm of Dimitriadis et al. (2017b), we also constructed networks using the NS, FA or MD as edge weights, due to their prevalent use in the literature (for example Honey et al. (2009) and Collin et al. (2016)), employing the OMST algorithm to select the edges in a data-driven manner.

#### 2.3.4 Thresholded graphs

Finally, we constructed graphs with topology determined by the CM that is weighted the NS, FA or MD, and a threshold applied to remove edges with the lowest weights. Instead of imposing an arbitrary threshold, the threshold was determined by imposing the constraint that the graphs exhibit the same sparsity as the OMST graphs that exhibited the highest reproducibility (more on this is included in Section 3). Once the topology of each of those graphs was specified, the weights of the edges were either kept as they were or re-weighted with one of the remaining two metrics.

#### 2.3.5 Summary of graphs investigated

The details of all the schemes that we considered are listed in Table 2 for easy reference. In the same table we also list the mean graph similarity (see Section 2.5) between scans for those schemes, for graphs generated from the DWIs with *b* = 2000 s/mm^2^, because that guided the decision of which schemes to look into in more detail. We discuss this more in Section 3.

**Table 2:**
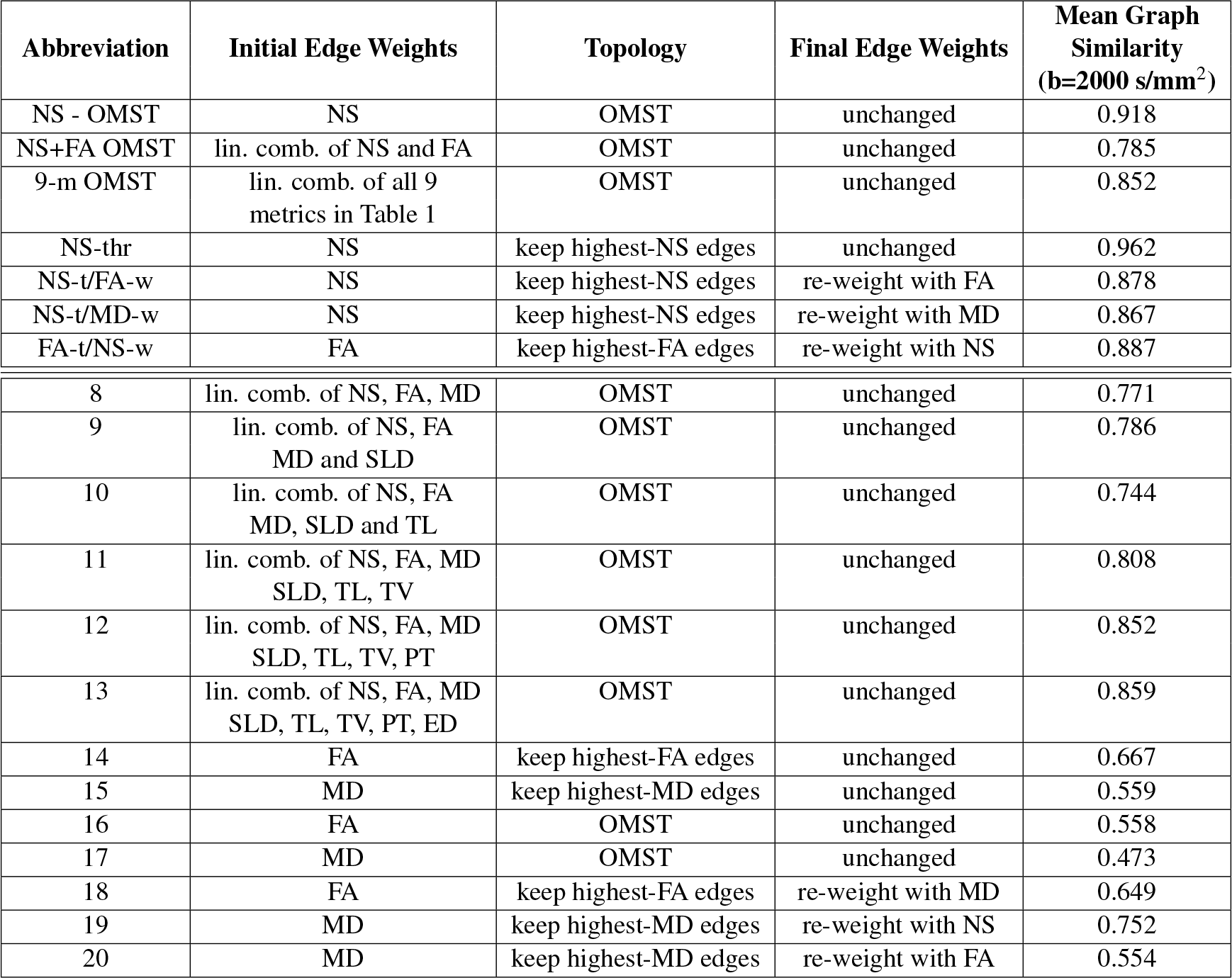
Graph-construction schemes. The first seven schemes are the ones that we discuss in detail in this article. The remaining 13 either give smaller reproducibility or are very similar to one of the first seven and therefore are not discussed in detail - and thus are numbered rather than given an abbreviation.

### 2.4 Graph Theory

We computed graph theoretical metrics for all the graphs using the Brain Connectivity Toolbox (Rubinov and Sporns, 2010). Specifically, we calculated the node degree, clustering coefficient, local efficiency and betweenness centrality for each node. We also calculated the global efficiency and the characteristic path length for each graph.

### 2.5 Comparison and Statistical Analysis

We performed three different forms of comparison:

1. Longitudinal Reproducibility: This was established by comparing the graphs from test and retest data, for each graph-construction scheme, independently for each diffusion weighting.
2. Between-Scheme Agreement: This was assessed by comparing graphs generated via different construction schemes, for each scan and diffusion weighting, for each participant.
3. Inter-B-Value Agreement: This was assessed by comparing graphs generated from the same scan using the same graph-construction scheme, but DWIs with two different diffusion weightings.

#### 2.5.1 Graph reproducibility

The *graph similarity* between two graphs was computed as the dot product of two vectors, each formed by concatenating the unique edges of each graph, normalized by the magnitudes of those two vectors. Even though it is informative, the similarity between two graphs encompasses both how similar the topologies of the graphs are (i.e. which edges exist in each graph) *and* how similar the edge weights are. In order to assess the impact of each of these contributions to the overall graph similarity, we looked into each contribution independently.

#### 2.5.2 Topology reproducibility

As a measure for the topology similarity, we constructed a vector containing as many entries as the total number of unique edges in our graphs, and assigned to each entry the value of 1 if the corresponding edge existed in both graphs being compared or if the edge did not exist in either of the graphs being compared, and the value of 0 if the edge in question existed in one of the two graphs being compared but not the other. We then averaged the entries of that vector to get the *topology similarity* between the two graphs.

#### 2.5.3 Edge reproducibility

In order to assess the edge reproducibility we calculated the intra-class correlation coefficient (ICC) for the edge weights, for all edges that appeared in both scans of at least 12 (one third) of the 37 participants in the group. The choice to use one third as a representative subgroup was partially arbitrary, but motivated by the fact that in order for the ICC to be a sensible measure, it is necessary to have a representative group of participants. We also calculated the absolute value of the fractional difference of the edge weights, namely the absolute value of the difference divided by the mean value of the edge weights, for each edge of each graph. The ICC gives the range of variability of each edge weight between scans in the context of the overall variability of the weights of that edge among participants. On the other hand, the absolute fractional difference gives a participant-specific measure of that variability.

We were also interested in whether dMRI-measured attributes of the WM tracts (for example mean FA, NS, etc.) are related to the reproducibility of edge weights. We constructed a general linear model (GLM) with the absolute fractional difference (between scans) of the edge weights as the dependent variable. We hypothesized that the fractional difference between edge weights would depend both on the attributes of the edges and on the absolute difference of those attributes between scans. Additionally, because the values of attributes of WM tracts are highly correlated between scans (a fact confirmed in our analysis, with all WM tract attributes exhibiting correlation coefficients of 0.83 or higher between scans, with *p*-values of 10^−7^ or lower), we used only the attributes listed in Table 1 for graphs generated from the first scan and their differences between scans, as independent variables (all variables were z-transformed).

We also wanted to assess whether the edges that appear in the graph from only one of the two scans have any particular attributes. We used two-sample t-tests to compare the distributions of the nine edge attributes for edges that appeared in both scans to their distributions for edges that appeared in only one of the two scans.

#### 2.5.4 Most significant edges

To identify the edges that have the highest weights across participants for each scheme, we selected the edges with weight of 0.9 or higher from the graphs of each participant and summed up the weights. The edges with the highest sums of edge weights were identified as the strongest for each scheme and each scan.

#### 2.5.5 Reproducibility of graph theoretical metrics

To assess the longitudinal reliability of graph theoretical metrics for the different graph-construction schemes, we calculated the ICC and the absolute value of the fractional difference for the graph theoretical metrics between the two scans, both for the local and the global graph theoretical metrics listed in Sec. 2.4.

### 2.6 Power of Structural Network Analyses

We were interested in evaluating the impact of different graph-construction schemes on the number of participants required in: a) comparative studies, namely studies where groups of participants exhibiting different characteristics (cognitive abilities, disease, etc.) are compared to each other; and b) longitudinal studies, namely studies in which specific measures are computed at two or more different time points for the same participants. A detailed calculation of the number of participants requires knowledge of the specific quantities of interest as well as of the populations involved, therefore such calculations need to be made on a case-by-case basis.

To estimate the impact of the choice of scheme on the power of such studies and on the required number of participants, we assume that we have two populations, each with the same number of participants *N*, and we measure the quantity *x* for each participant. Alternatively, we have a population with *N* participants who undergo an intervention, and we measure the quantity *x* for all participants before and after the intervention. We also assume that the distribution *p*_1_(*x*) of *x* for the first population (in the former scenario) or for the population before the intervention (in the latter scenario) is normal with a mean of 0, while for the second population (in the former scenario) or for the population after the intervention (in the latter scenario) the distribution *p*_2_(*x*) is a normal with a mean of *µ*. Finally, we assume that the standard deviations (SD) of both distributions are equal to each other, and denoted by *σ*. As demonstrated by Mumford (2012), if we specify the confidence level at which we wish to detect a possible difference in the means of the distributions, the quantity

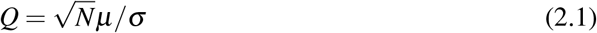

determines the power of the study to identify a possible effect. The higher *Q* is, the higher the power of the study. This means that the tighter the distributions (lower *σ*), the lower the number of participants needed to detect the effect. The fact that it is the square root of *N* that shows up in *Q* means that even a modest improvement in *σ* can lead to a significant reduction in the number of participants required to detect a given effect with the same power, for a given statistical-significance threshold. On the other hand, an increase in *σ* can lead to a deterioration of the power of a study, if not enough participants can be recruited to allow for confident detection of a given effect. If the number of participants cannot be changed, the dependence of *Q* on the mean *µ* of the second distribution indicates that reducing *σ* implies that a smaller value of *µ*, and thus of deviation from the zero-mean distribution, is required for the effect to be observed with a given power at a given statistical-significance threshold. Taking this into account, in order to assess the impact of the choice of graph-construction scheme of the various metrics of interest, we calculated the SD of their distributions. We then calculated the increase in the number of participants that would be required, for all the graph-construction schemes in comparison to the scheme with the lowest SD for the metrics of interest.

### 2.7 A Note on b-Values

As mentioned in Sec. 2.1, the DWIs were acquired for three different diffusion weightings. *B*-values of up to 2000 s/mm^2^ give very good fit for the diffusion tensor (DT) and its fractional anisotropy, while for higher *b*-values the higher order effects of diffusion need to be taken into account and kurtosis terms need to be included in the fit (Jensen et al., 2005; Jensen and Helpern, 2010; Tabesh et al., 2011). On the other hand, *b*-values of under 1500 s/mm^2^ can be problematic when it comes to resolving crossing fibers with CSD tractography algorithms (Tuch et al., 2002; Tournier et al., 2004, 2008; Cho et al., 2008), which is important given the prevalence of such fibers in the human brain (Jeurissen et al., 2012). Here, we present the results derived from DWIs acquired with *b* = 2000 s/mm^2^ in more detail than for the other weightings, because, unlike the other two diffusion weightings, this one gives reliable tractography results with the CSD algorithm *and* reliable results for the DT fits. It is, however, important to understand how similar or different the results are for the other two diffusion weightings in comparison to *b* = 2000 s/mm^2^, specifically for single-shell experiments or datasets as will be discussed in Sec. 4. We therefore also present the reproducibility results for those two diffusion weightings.

## 3 Results

As explained in Sec. 2, we considered twenty graph-construction schemes (see Table 2). We present detailed results for the seven schemes that a) resulted in reproducibility (for the graphs and their various attributes) that is high enough for the graphs to be useful for comparative and longitudinal studies and b) are different enough from each other to convey a different picture of the structural connectome. Specifically, we excluded all schemes that resulted in mean graph similarity over participants of under 0.75 (schemes 10, 14, 15, 16, 17, 18 and 20 in Table 2; also see Messaritaki et al. (2019) for more details on the graph similarity for the FA- and MS-schemes). We also excluded schemes for which the graph similarity was under 0.7 for any of the participants regardless of the mean graph similarity (schemes 8, 9, 19 in Table 2). Schemes 11, 12 and 13 resulted in graphs that are very similar to the 9-m OMST scheme, and we decided to discuss the latter in detail due to the fact that it exhibits higher reproducibility for the graph theoretical metrics. The schemes we discuss in detail, then, and the abbreviations used to refer to them, are:

- Scheme 1: The edge weights are equal to the NS, and the OMST algorithm is used to select the edges (NS-OMST).
- Scheme 2: The edge weights are equal to a linear combination of the NS and the FA, with the coefficients identified by the diffusion-distance algorithm. The OMST algorithm is used to select the edges (NS+FS OMST). This is the same scheme that is described by Dimitriadis et al. (2017b), using only two metrics to weigh the edges.
- Scheme 3: The edges are weighted by linear combinations of 9 metrics, with the coefficients identified via the diffusion-distance algorithm. The OMST algorithm is used to select the edges (9-m OMST).
- Scheme 4: The edges are weighted by the NS, and an absolute threshold is used to keep the edges that have the highest NS (NS-thr).
- Scheme 5: The edges are first weighted by the NS and thresholded to keep the ones with the highest NS. Then they are re-weighted by their FA. In other words, the topology of the graph is defined by the thresholded-NS scheme and the weights are equal to the FA (NS-t/FA-w).
- Scheme 6: The edges are first weighted by the NS and thresholded to keep the ones with the highest NS. Then they are re-weighted by their MD. In other words, the topology of the graph is defined by the thresholded-NS scheme and the weights are equal to the MD (NS-t/MD-w).
- Scheme 7: The edges are first weighted by the FA and thresholded to keep the ones with the highest FA. Then they are re-weighted by their NS. In other words, the topology of the graph is defined by the thresholded-FA scheme and the weights are equal to the NS (FA-t/NS-w).

We remind the reader that, for the four thresholded schemes, the threshold was defined so that the networks exhibited the same sparsity as the corresponding 9-m OMST networks, which had the highest mean network similarity of the OMST networks that use more than one metric (see Sec. 3.2.1).

### 3.1 Representative Graphs

Fig. 1 shows the graphs for the structural networks of the first scan of one participant for the seven different schemes, derived from DWIs acquired with *b* = 2000 s/mm^2^.

**Figure 1:**
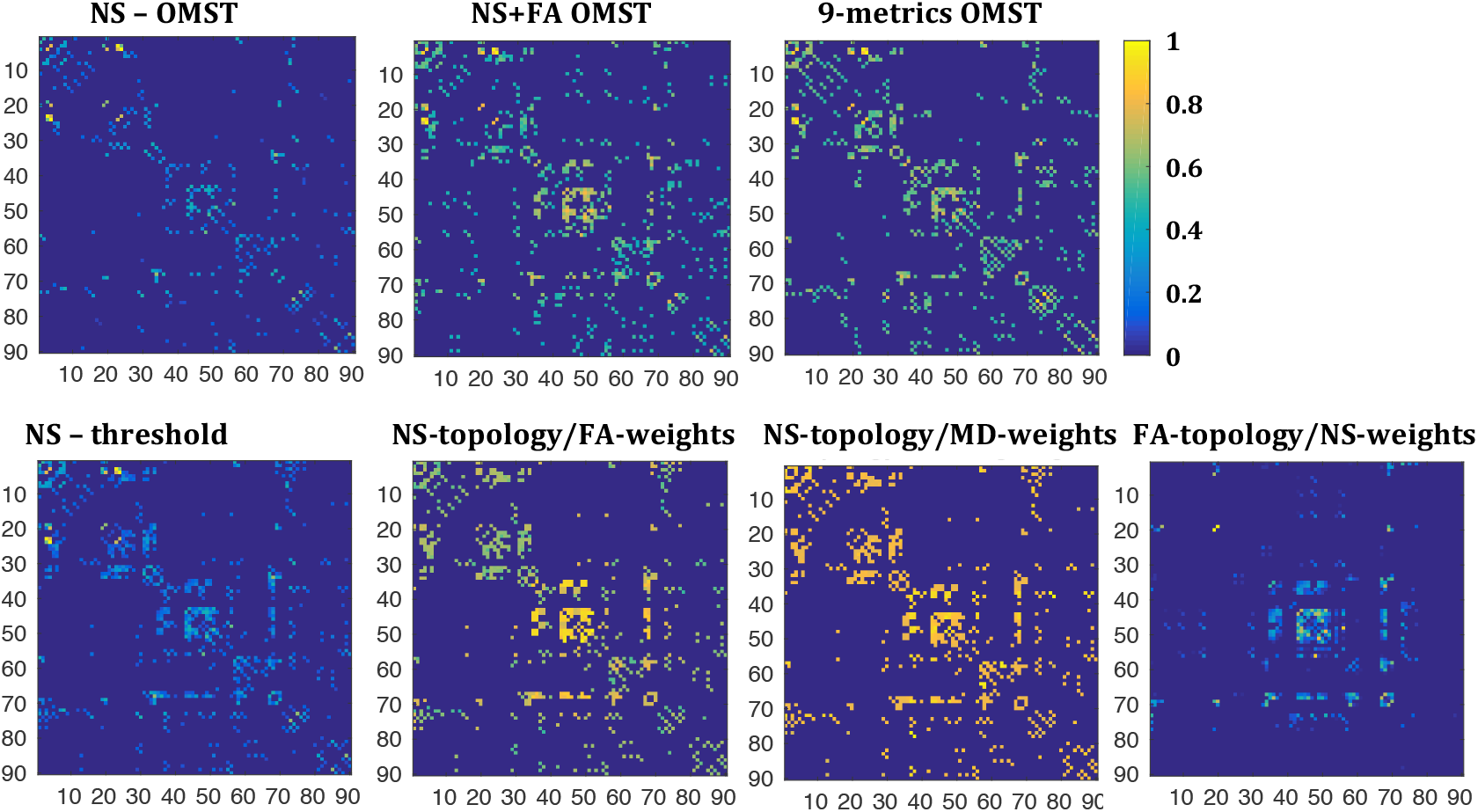
Sample graphs representing the structural connectome for the first scan of one of the participants, derived from the DWIs with *b* = 2000 s/mm^2^. All graphs have been normalized so that the maximum edge weight within each graph is 1. The colors in the colorbar correspond to the edge weights. By construction, the NS-OMST graph is the sparsest, while the other graphs have the same sparsity as each other.

For six of the seven graph-construction schemes the majority of the edges close to the diagonal were non-zero. The graph resulting from the FA-t/NS-w scheme was the only exception, in which the edges close to the diagonal for nodes up to node 30 were not present, or not as strong in relation to the rest of the edges as for the other schemes.

The weights of the edges were very variable between schemes, as was the relative weight of the edges within each scheme. For the NS-OMST and NS-thr schemes, which rely exclusively on the NS for the topology and edge weights, only a few edges had weights close to 1, while all the others had weights of 0.6 or lower. For the NS-t/MD-w graph, on the other hand, most edges had values close to 1, reflecting the fact that the MD of white matter tracts is quite uniform in the human brain. The remaining four schemes exhibited a more uniform distribution of edge weights between the values of 0 and 1.

#### 3.1.1 Between-scheme similarity of graphs

The mean of the graph similarity between schemes (over the 37 participants and the two scans) is given in Table 3 for all pairs of graph-construction schemes. The high similarity of the graphs generated by the NS-t/FA-w and NS-t/MD-w schemes is, of course, a result of their topologies being identical. The graphs generated by the NS-OMST and NS-thr schemes were also highly similar to each other, which is due to the fact that the weights used for the edges are the same. Notably, this is true despite of the fact that these graphs generally exhibited very different sparsities (the mean sparsity of the NS-OMST graphs across participants was 0.044 while that of the NS-thr graphs was 0.096.)

**Table 3:** Mean between-scheme similarity of graphs, for *b* = 2000 s/mm^2^. The mean is over all participants and then over the two scans. The highest values of the similarity are indicated in bold letters.

|  |  |  |  |  |  |  |
| --- | --- | --- | --- | --- | --- | --- |
| **NS+FA OMST** | 0.633 |  |  |  |  |  |
| **9-m OMST** | 0.637 | 0.617 |  |  |  |  |
| **NS-thr** | **0.883** | 0.754 | 0.744 |  |  |  |
| **NS-t/FA-w** | 0.538 | 0.696 | 0.685 | **0.808** |  |  |
| **NS-t/MD-w** | 0.537 | 0.664 | 0.694 | 0.798 | **0.988** |  |
| **FA-t/NS-w** | 0.552 | 0.539 | 0.427 | 0.637 | 0.503 | 0.438 |
|  | **NS OMST** | **NS+FA OMST** | **9-m OMST** | **NS-thr** | **NS-t/FA-w** | **NS-t/MD-w** |

### 3.2 Reproducibility Results

#### 3.2.1 Graph reproducibility

The between-scan graph similarity for the 37 participants is shown in Fig. 2. We show the similarity for the graphs constructed via the OMST schemes separately from that for the four thresholded schemes, using the same scale on the vertical axis in both plots.

**Figure 2:**
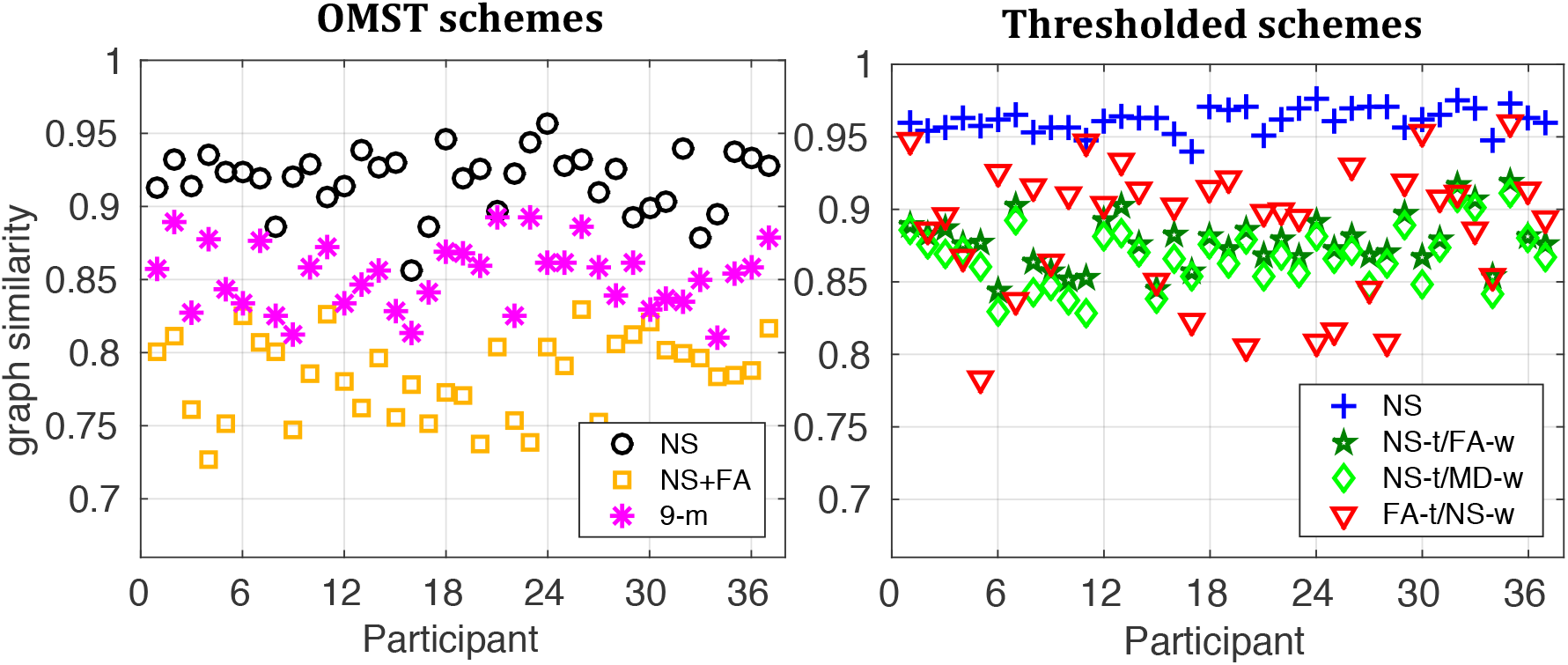
Graph similarity between the two scans for the 37 participants, for *b* = 2000 s/mm^2^. The similarity for the OMST schemes is plotted separately from that for the thresholded schemes, however both plots use the same scale for ease of comparison.

The NS-OMST had the highest graph similarity among the OMST schemes, just over 0.9 for most participants. Of the seven schemes considered, the NS-thresholded scheme resulted in the highest similarity, about 0.95 for all participants. Notably, even though the FA-t/NS-w scheme resulted in a very high mean graph similarity, it also resulted in the widest range of graph similarity among the 37 participants, ranging from 0.75 to 0.95.

The mean and SD of the graph similarity for the three different diffusion weightings is given are Table 4. The values of the mean similarity were very close to each other for the graphs generated with the same scheme across *b*-values, for all seven schemes. We compared the distributions of the graph similarity for each scheme for all pairs of the three diffusion weightings with paired t-tests, and applied false-discovery-rate (FDR) multiple comparison correction with a threshold of *p <* 0.01. The results are shown in Table 5, where “X” indicates pairs of distributions that are statistically significantly different from each other. There are no statistically significant differences in the graph similarity distributions between *b* = 2000 s/mm^2^ and *b* = 3000 s/mm^2^ for any of the seven schemes, indicating that the graph similarities are not affected by the *b*-value for those stronger diffusion weightings. Additionally, the 9-m OMST scheme does not show any statistically significant differences in the similarity distributions regardless of the pair of *b*-values compared. It is possible that using several metrics to weigh the edges reduces the effect of inaccuracies in the evaluation of tract metrics, and therefore makes that scheme more robust across *b*-values, as far as the graph similarity is concerned.

**Table 4:** Mean and SD of the similarity distributions of the graphs for the three diffusion weightings.

|  |  | NS OMST | NS+FA | 9-m OMST | NS-thr | NS-t/FA-w | NS-t/MD-w | FA-t/NS-w |
| --- | --- | --- | --- | --- | --- | --- | --- | --- |
| b: 1000 s/mm<sup>2</sup> | mean | 0.901 | 0.763 | 0.852 | 0.952 | 0.862 | 0.845 | 0.855 |
|  | SD | 0.020 | 0.034 | 0.026 | 0.010 | 0.022 | 0.023 | 0.045 |
| b: 2000 s/mm<sup>2</sup> | mean | 0.918 | 0.785 | 0.852 | 0.962 | 0.878 | 0.867 | 0.888 |
|  | SD | 0.021 | 0.028 | 0.023 | 0.008 | 0.018 | 0.020 | 0.041 |
| b: 3000 s/mm<sup>2</sup> | mean | 0.917 | 0.779 | 0.847 | 0.923 | 0.878 | 0.865 | 0.882 |
|  | SD | 0.023 | 0.025 | 0.025 | 0.009 | 0.021 | 0.023 | 0.051 |

**Table 5:** Comparison of similarity distributions across *b*-values (paired t-tests). The cells marked with X indicate the pairs of *b*-values for which the distributions were statistically significantly different, after FDR correction for multiple comparisons has been applied, at a *p*-value threshold of 0.01.

|  | 1000 s/mm<sup>2</sup> vs. 2000 s/mm<sup>2</sup> | 1000 s/mm<sup>2</sup> vs. 3000 s/mm<sup>2</sup> | 2000 s/mm<sup>2</sup> vs. 3000 s/mm<sup>2</sup> |
| --- | --- | --- | --- |
| NS OMST | X | X |  |
| NS+FS OMST | X |  |  |
| 9-m OMST |  |  |  |
| NS-thr | X | X |  |
| NS-t/FA-w | X | X |  |
| NS-t/MD-w | X | X |  |
| FA-t/NS-w | X |  |  |

We also compared the distributions of graph similarity for graphs generated with two different schemes separately for each diffusion weighting, for all possible pairs of schemes (paired t-test, after FDR multiple comparison corrections at a threshold of *p* = 0.01). The results are shown in Table 6. Blank cells indicate pairs of schemes for which the distributions are statistically significantly different for all three *b*-values. The numbers in the filled cells indicate the *b*-values for which the graph similarity distributions are not statistically significantly different. For most of the pairs of schemes, the graph similarity distributions are statistically significantly different from each other, indicating that the graph similarity depends on the scheme. For five pairs of schemes, the distributions were not statistically significantly different for *b* = 1000 s/mm^2^, while two of these pairs of schemes also resulted in distributions that were not statistically significantly different for the two higher diffusion weightings.

**Table 6:** Comparison of distributions of network similarity across schemes, for the 3 *b*-values (paired t-tests, followed by FDR multiple-comparison correction with *p*-value threshold of 0.01). Blank cells indicate pairs of schemes for which the distributions are statistically significantly different for all three *b*-values. The numbers in the filled cells indicate the *b*-values (in s/mm^2^) for which the similarity distributions are not statistically significantly different.

|  |  |  |  |  |  |  |
| --- | --- | --- | --- | --- | --- | --- |
| **NS+FA** |  |  |  |  |  |  |
| **9-m OMST** |  |  |  |  |  |  |
| **NS-thr** |  |  |  |  |  |  |
| **NS-t/FA-w** |  |  | 1000 |  |  |  |
| **NS-t/MD-w** |  |  | 1000 |  |  |  |
| **FA-t/NS-w** |  |  | 1000 |  | 1000, 2000, 3000 | 1000, 2000, 3000 |
|  | **NS OMST** | **NS+FA** | **9-m OMST** | **NS-thr** | **NS-t/FA-w** | **NS-t/MD-w** |

#### 3.2.2 Topology reproducibility

The topology similarity is shown in Fig. 3 for the different schemes for *b* = 2000 s/mm^2^. The mean and the SD of the topology similarity for each scheme are given in Table 7, for the three diffusion weightings. The topology similarity was over 0.9, for all participants regardless of the scheme used to construct the graphs. The NS-OMST scheme had the highest mean and the smallest SD of topology similarity for all three diffusion weightings. The NS-thr, NS-t/FA-w and NS-t/MD-w schemes all had the same topology similarity since they share the same topology.

**Table 7:** Mean and SD of the topology similarity distributions of the graphs for the 3 *b*-values.

|  |  | NS OMST | NS+FA | 9-m OMST | NS-thr | NS-t/FA-w | NS-t/MD-w | FA-t/NS-w |
| --- | --- | --- | --- | --- | --- | --- | --- | --- |
| b: 1000 s/mm<sup>2</sup> | mean | 0.981 | 0.944 | 0.966 | 0.973 | 0.973 | 0.973 | 0.931 |
|  | SD | 0.003 | 0.008 | 0.005 | 0.004 | 0.004 | 0.004 | 0.009 |
| b: 2000 s/mm<sup>2</sup> | mean | 0.984 | 0.947 | 0.963 | 0.974 | 0.974 | 0.974 | 0.931 |
|  | SD | 0.002 | 0.007 | 0.007 | 0.005 | 0.005 | 0.005 | 0.011 |
| b: 3000 s/mm<sup>2</sup> | mean | 0.983 | 0.947 | 0.962 | 0.974 | 0.974 | 0.974 | 0.934 |
|  | SD | 0.003 | 0.005 | 0.006 | 0.005 | 0.005 | 0.005 | 0.010 |

**Figure 3:**
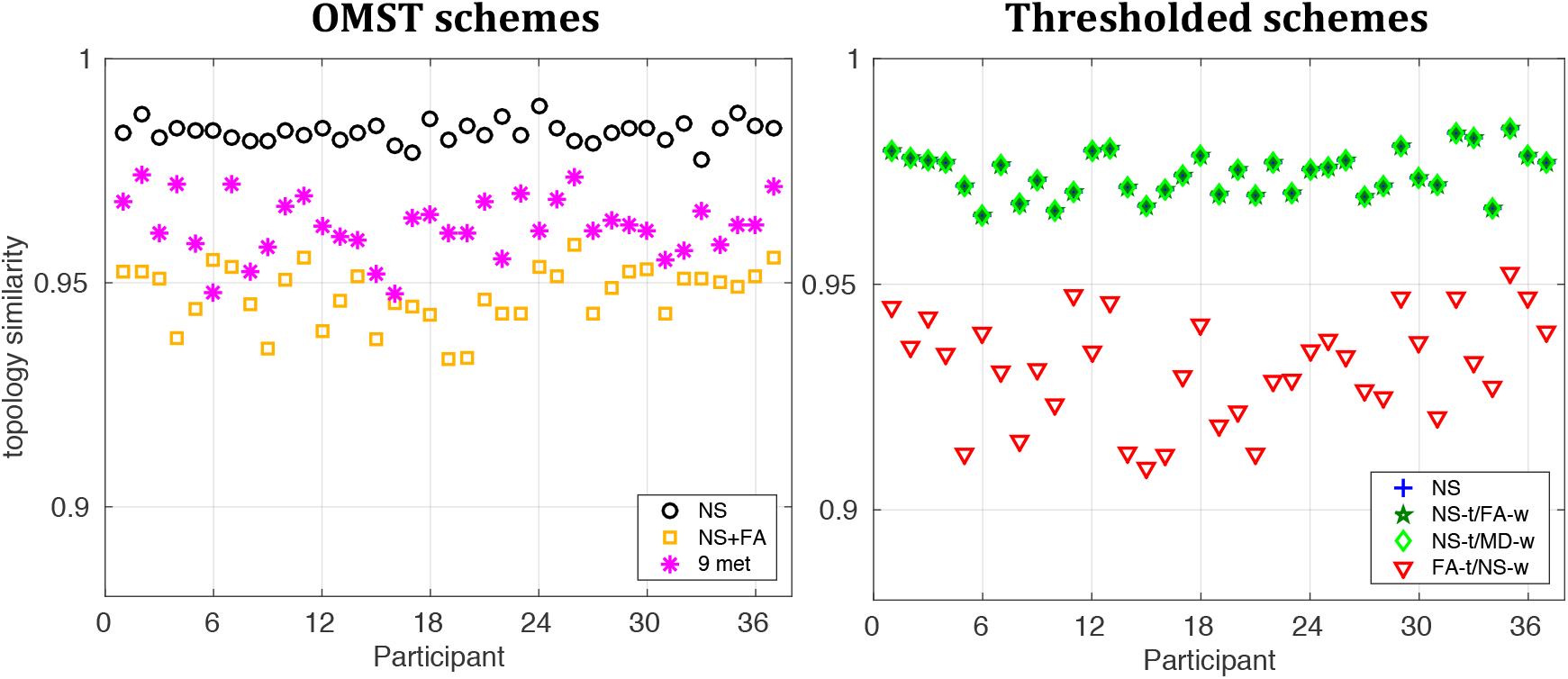
Topology similarity between the two scans for the 37 participants, for *b* = 2000 s/mm^2^. For clarity of the plot, the similarity for the OMST schemes is plotted separately from that for the thresholded schemes, however both plots use the same scale for ease of comparison.

We compared the distributions of the topology similarity for the five schemes (omitting the NS-t/FA-w and NS-t/MD-w because they have the same topology as the NS-thr scheme) with paired t-tests, for each diffusion weighting. For all three diffusion weightings, all pairs of schemes exhibited statistically significantly different distributions of topology similarity (all *p* < 10^−6^, and survived FDR multiple comparison corrections for a *p*-value threshold of 0.01), indicating that the topology similarity is dependent on the chosen scheme.

#### 3.2.3 Edge reproducibility

The ICC for the edge weights is shown in Fig. 4, for graphs generated from the DWIs acquired with *b* = 2000 s/mm^2^. For all except the 9-m OMST scheme, the majority of the edge weights exhibited ICCs above 0.7. In order for the plots in Fig. 4 to be easily interpretable, the scale was set to be between 0 and 1 and therefore the few edges that have ICCs less than zero, i.e., edges for which the between-scan variability of the edge-weights is larger than the range of values of those weights, are not evident. Fig. 5 shows the percentage of edges versus ICC for the seven schemes, with the full range of values for each scheme. The FA-t/NS-w scheme resulted in the highest percentage, 66.4%, of edges with ICCs of 0.85 or higher, while the NS-thr and NS-t/FA-w schemes had 61.9% and 61.4% edges with ICCs greater than 0.85 respectively. It is noteworthy that the 9-m OMST schemes resulted in distributions that are shifted to lower values compared to those of the other six schemes. That is possibly due to the fact that the nine WM tract metrics that are combined to form the edge weights result in some added variability that leads to larger ICCs.

**Figure 4:**
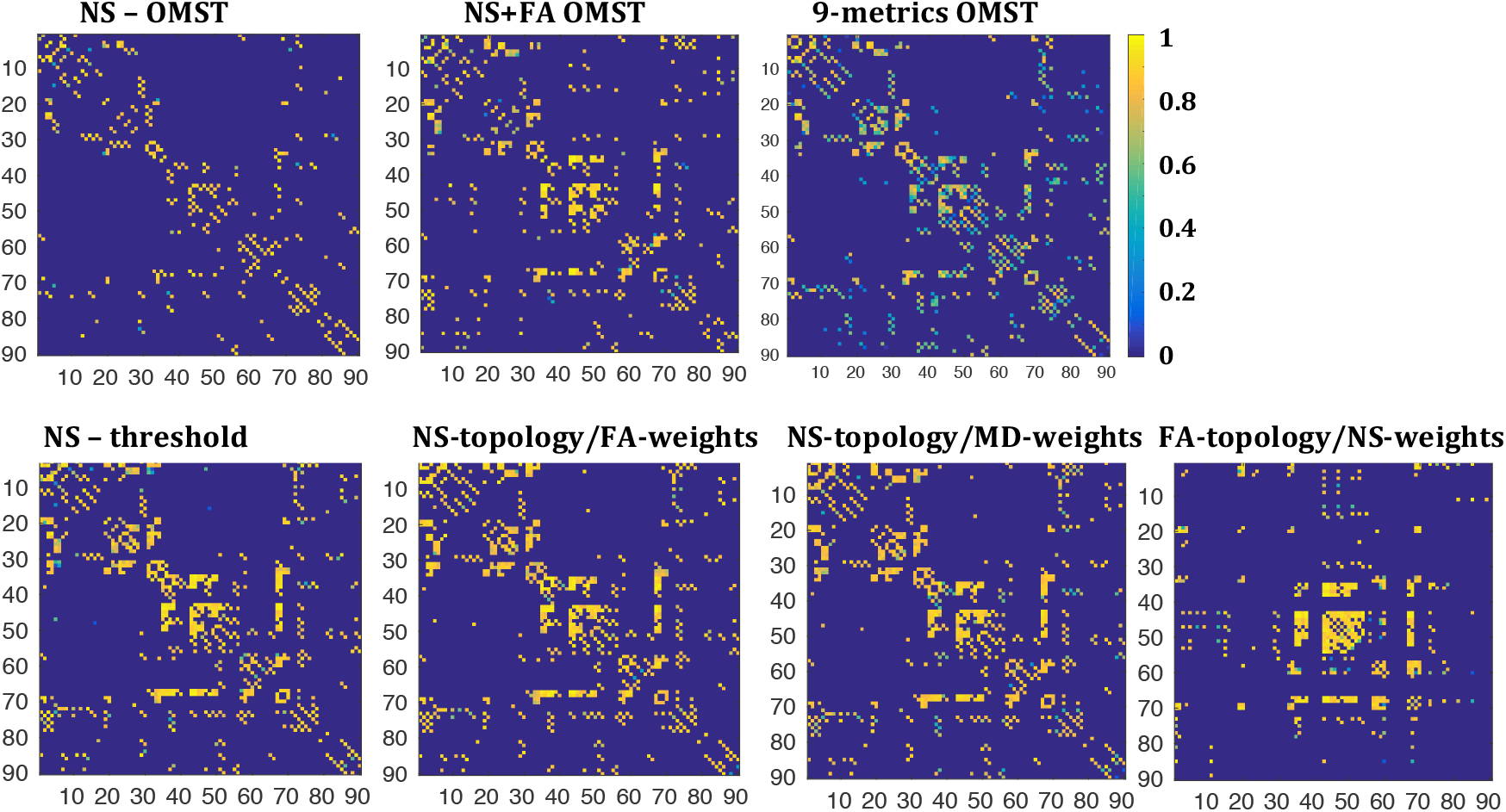
ICC of the edges present in both scans of at least 12 of the 37 participants, for the different schemes considered.

**Figure 5:**
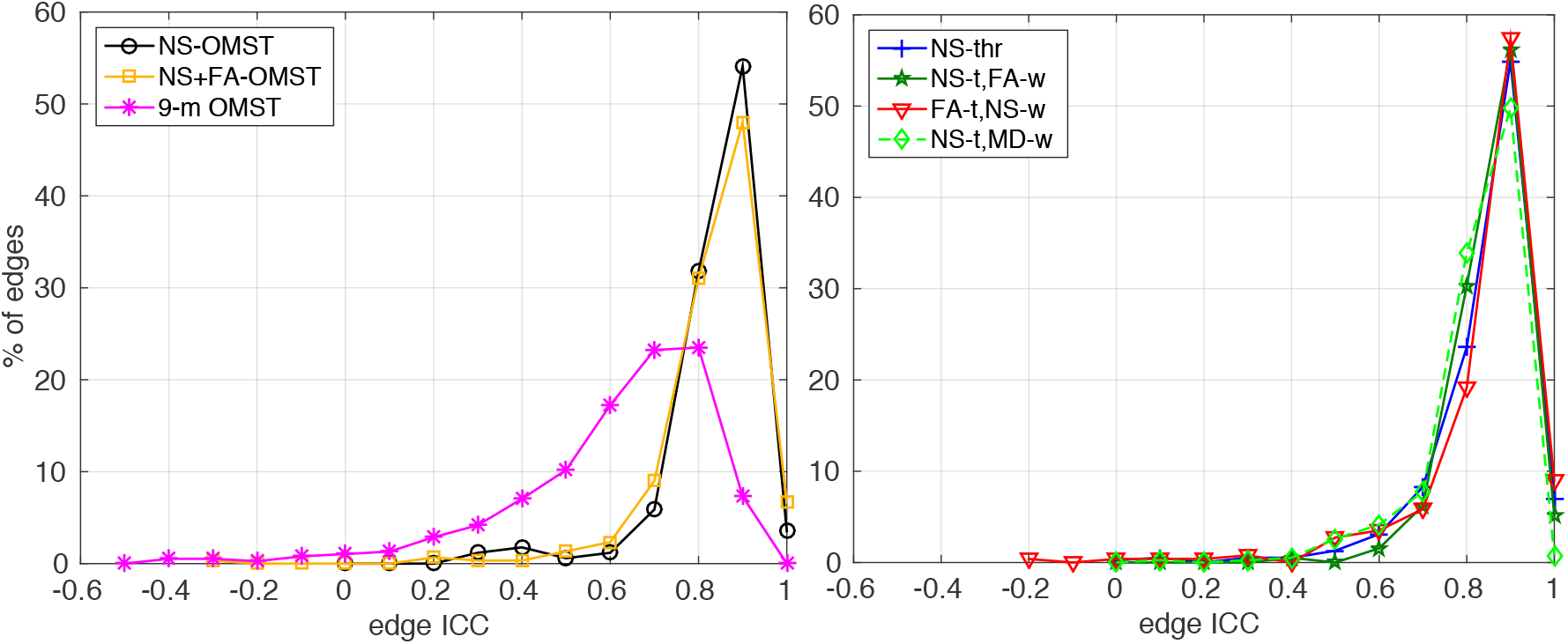
Percentage of edges versus ICC, for *b* = 2000 s/mm^2^. Left panel: OMST schemes, right panel: thresholded schemes. Only edges that show up in the graphs from both scans of at least 12 participants have been included.

Fig. 6 shows the percentage of edges in the graphs of all participants versus the absolute fractional difference between scans of the edge weights, for graphs generated from the DWIs acquired with *b* = 2000 s/mm^2^. In each panel, the points at the left part of the plot correspond to edges for which the absolute fractional difference is small and therefore the weights do not differ much between the two scans. The points at the right part of the plot correspond to edges that appear in one of the two scans of a given participant, but not the other, and as a result have absolute fractional differences equal to 2. For a scheme to be deemed reliable as far as the reproducibility of edges and edge-weights is concerned, it must result in most of the edges having small edge-weight fractional differences, and zero, or as few as possible, edges with an absolute fractional difference of edge weights equal to 2.

**Figure 6:**
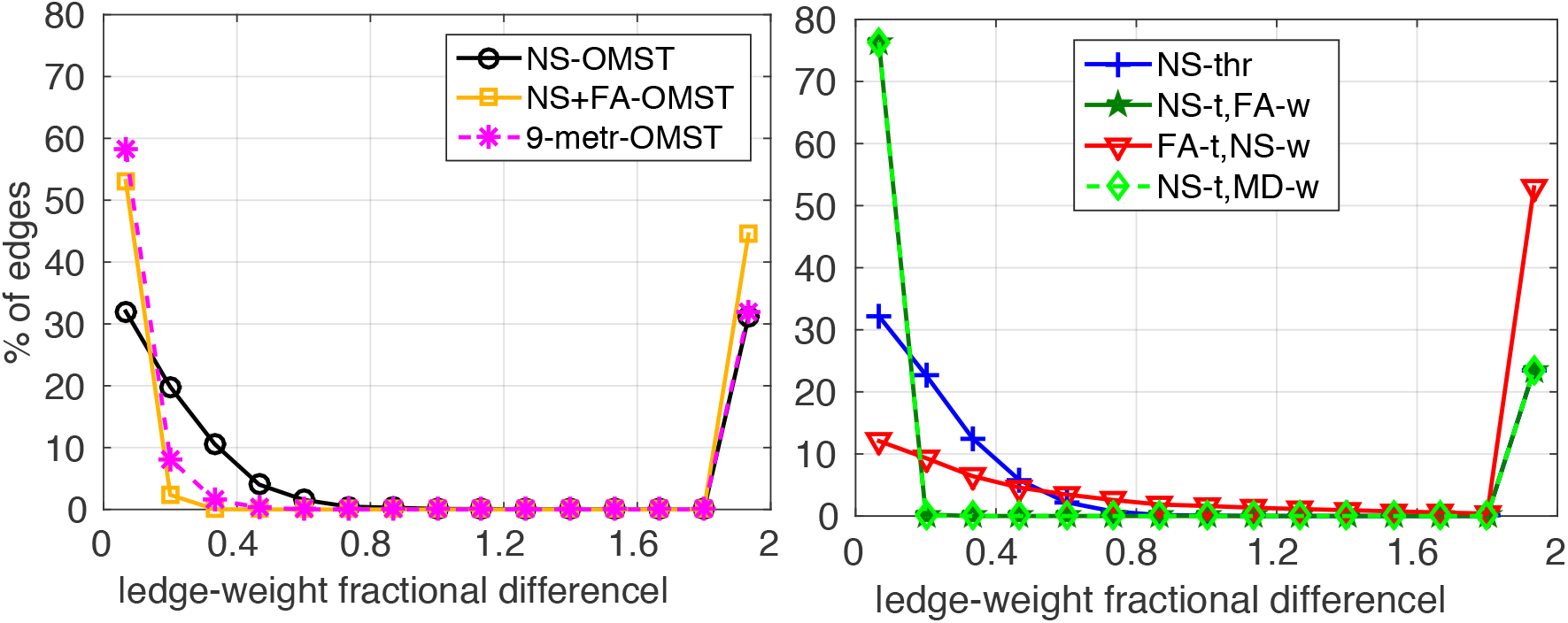
Percentage of edges versus absolute fractional difference in the edge weights in the graphs from the two scans, for *b* = 2000 s/mm^2^. Left panel: OMST schemes, right panel: thresholded schemes. The points at the rightmost side of each figure correspond to absolute fractional difference of 2, namely to edges that appear in graphs from only one of the two scans.

The NS-t/FA-w and NS-t/MD-w schemes performed the best, with 76.2% of the edges exhibiting very small fractional differences and only 23.6% of the edges appearing in the graphs from only one of the two scans. The NS-thr scheme resulted in the same small percentage of edges appearing in the graphs from only one of the two scans, but it also resulted in 75.4% of the edges exhibiting fractional differences of up to 0.6 in the edge weights. Of the OMST schemes, the 9-metric scheme was the best because it resulted in nearly 68% of the edges having a fractional difference of under 0.33, while 31.9% of the edges appeared in the graphs from only one of the two scans. We note that these percentages are correct for the graphs generated from the DWIs with *b* = 2000 s/mm^2^. However, the conclusions about which schemes do better also hold for the graphs generated from the DWIs with *b* = 1000 s/mm^2^ and *b* = 3000 s/mm^2^, and the distributions shown in Figures 5 and 6 were very similar for those two *b*-values.

The results of the GLM analysis (see Sec. 2) are given in Table 8. For the NS-OMST, NS+FA OMST and NS-thr schemes, the absolute difference in the number of streamlines of the edge between the two scans was the most significant predictor variable, and the larger the absolute difference in NS, the larger the absolute fractional difference in edge weights. For the NS-OMST scheme, the tract volume explained some of the variability, with the negative regression coefficient indicating that the smaller the volume of a tract, the larger the absolute fractional difference in edge weights. For the NS-thr scheme, the number of streamlines in the first scan was also significantly and negatively correlated with the absolute fractional difference in edge weight. In other words, edges that have a small number of streamlines are less reliably reproduced. For the NS-t/FA-w and NS-t/MD-w schemes, the absolute difference (between scans) in the metric used to weigh the edges was the best predictor, and positively correlated with the edge-weight absolute fractional difference. Finally, and somewhat surprisingly, for the FA-t/NS-w scheme, the best predictor variable was the TV in the first scan, with the absolute difference in TV between scans also contributing. For the 9-m OMST scheme, only a small fraction of the variability was explained by the attributes of the edges, and the coefficients of the GLM, even though statistically significant, were very small (less than 0.1) and therefore not useful in predicting which edges are likely to be highly reproducible. That could be due to the fact that the linear combination of metrics used to weigh the edges in the graphs is different among participants.

**Table 8:** GLM for the absolute value of the fractional difference of edge weights, for edges that appear in both scans of a given participant. The *p*-values for all the coefficients shown here were smaller than 10^−50^, and they all survived FDR multiple comparison correction. The variables are listed in order of significance.

|  | NS OMST | NS+FA OMST | NS-thr | NS-t/FA-w | NS-t/MD-w | FA-t/NS-w |
| --- | --- | --- | --- | --- | --- | --- |
| **variables** | $\vert \Delta(\text{NS})\vert : 0.63$<br>$\text{TV}_1: -0.50$ | $\vert \Delta(\text{NS})\vert : 0.55$<br>$\vert \Delta(\text{FA})\vert : 0.22$ | $\vert \Delta(\text{NS})\vert : 0.73$<br>$\text{NS}_1: -0.46$ | $\vert \Delta(\text{FA})\vert : 0.71$<br>$\text{FA}_1: -0.22$ | $\vert \Delta(\text{MD})\vert : 0.41$ | $\text{TV}_1: -0.60$<br>$\vert \Delta(\text{TV})\vert : 0.39$ |
| **% variability explained** | 39.2 | 33.2 | 48.1 | 57.2 | 16.5 | 31.7 |

The comparison of the distributions of edge attributes between edges that appear in graphs from both scans and those that appear in graphs from only one of the two scans showed no statistically significant differences. However, the edges that appear in graphs from only one of the two scans generally had low NS and low TV, for all graph-construction schemes except the FA-t/NS-w. This is shown in Figures 7 and 8, where it is also evident that for the NS-OMST, NS-thr and FA-t/NS-w schemes, edges with low NS but large differences in the NS between scans exhibited the largest absolute fractional differences in edge weights.

**Figure 7:**
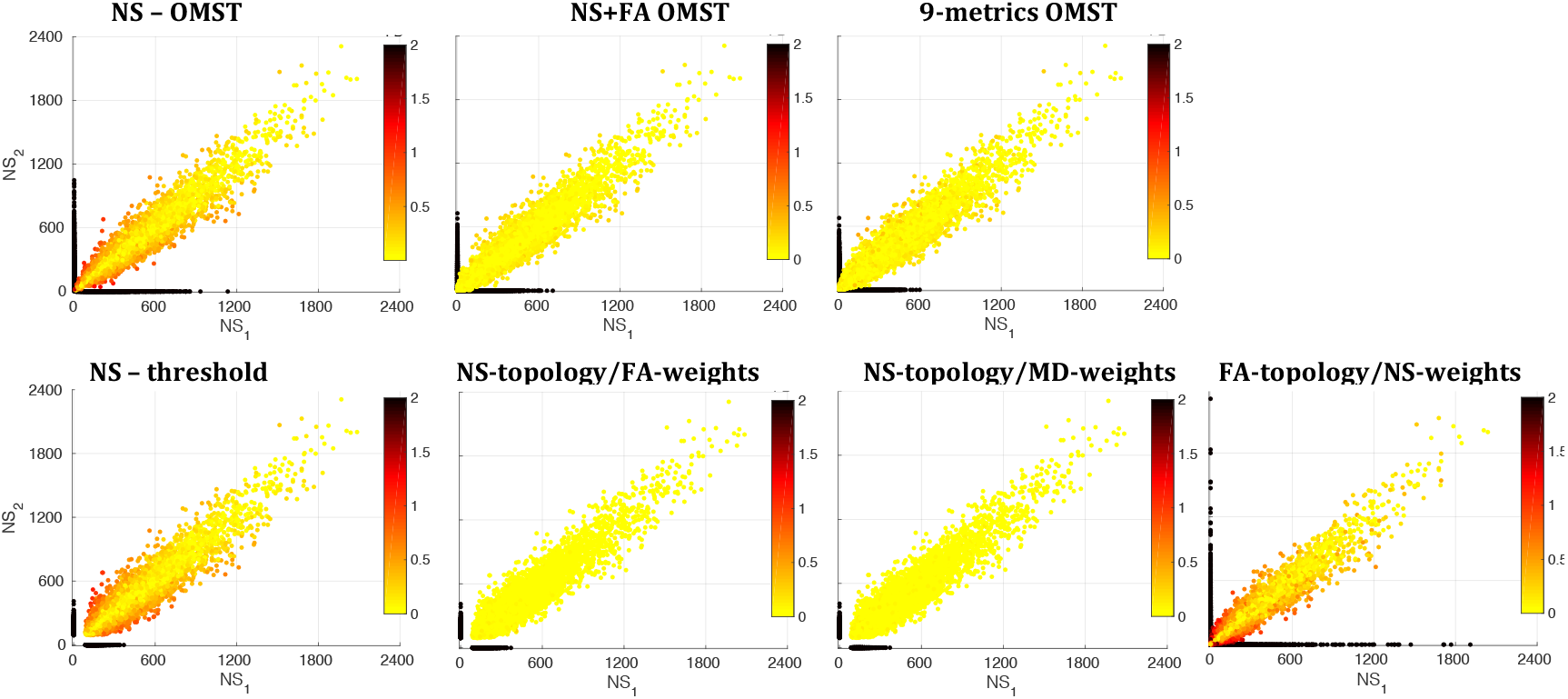
NS in scan 1 versus NS in scan 2, color-coded by the absolute value of the fractional difference in edge weights. The black dots correspond to edges that appear in the graphs from one of the two scans but not the other, and therefore have NS equal 0 in the scan in which the edge is not reproduced.

**Figure 8:**
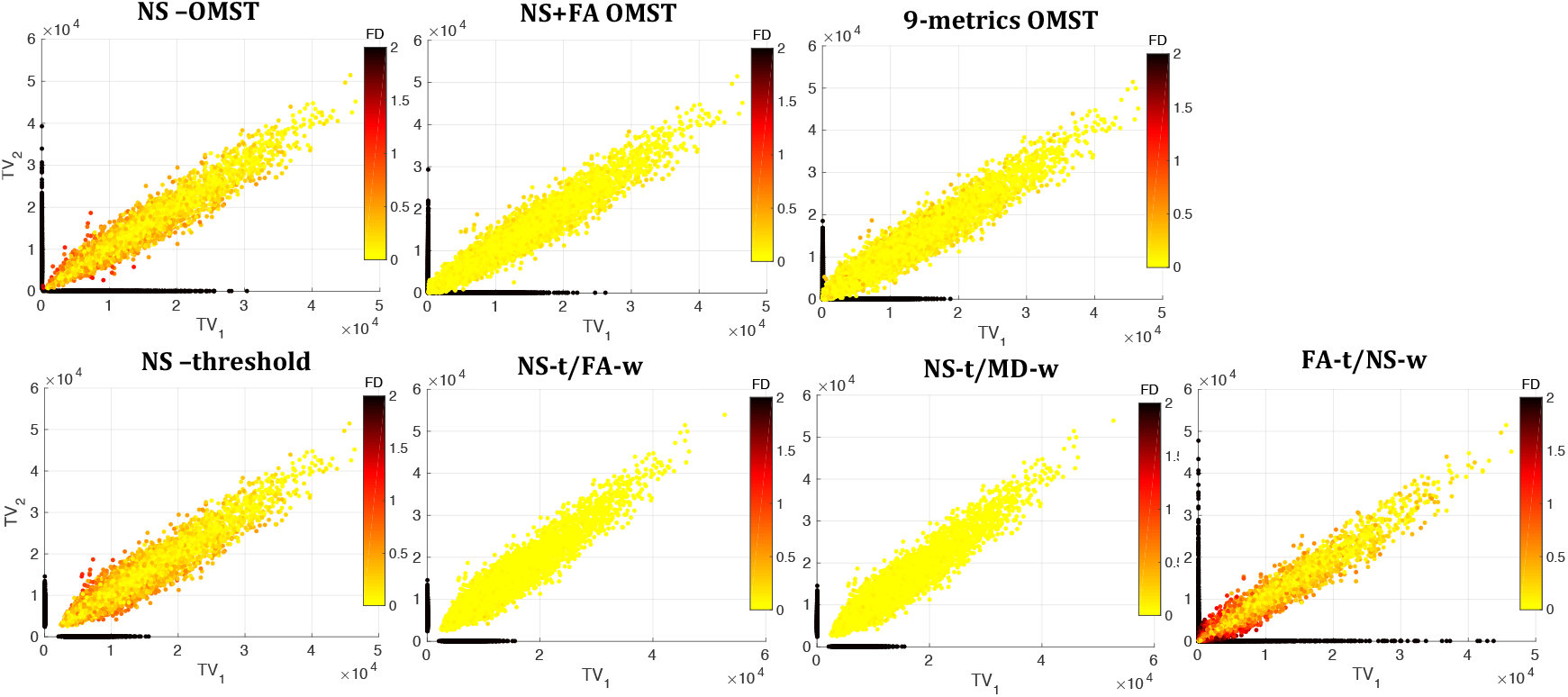
TV in scan 1 versus TV in scan 2, color-coded by the absolute value of the fractional difference in edge weights. The black dots correspond to edges that appear in the graphs from one of the two scans but not the other, and therefore have TV equal to 0 in the scan in which the edge is not reproduced.

#### 3.2.4 Strongest edges

The edges that are the strongest in all participants depended on the scheme used. The AAL regions interconnected by those edges are listed in Table 9 for the seven graph construction schemes.

**Table 9:** Connections that are strongest for all participants for the seven graph construction schemes.

| **Areas Connected (AAL atlas)** | **Schemes** |
| --- | --- |
| L superior frontal gyrus - L medial frontal gyrus | NS-OMST; (NS+FA)-OMST<br>9-m OMST; NS-thr |
| R superior frontal gyrus - R medial frontal gyrus | NS-OMST; (NS+FA)-OMST<br>9-m OMST; NS-thr |
| L supplementary motor area - R supplementary motor area | (NS+FA)-OMST; 9-m OMST; FA-t/NS-w |
| L hippocampus - L thalamus | NS-t/MD-w |
| R hippocampus - R thalamus | NS-t/MD-w |
| L hippocampus - R hippocampus | NS-t/MD-w |
| L hippocampus - L amygdala | NS-t/MD-w |
| R hippocampus - R amygdala | NS-t/MD-w |
| L putamen - L globus pallidus | NS-thr |
| R putamen - R globus pallidus | NS-thr |
| L posterior cingulate gyrus - L superior occipital gyrus | NS-t/FA-w |
| L posterior cingulate gyrus - R superior occipital gyrus | NS-t/FA-w |
| R posterior cingulate gyrus - L superior occipital gyrus | NS-t/FA-w |
| L cuneus - R cuneus | NS-t/FA-w |
| R calcarine sulcus - L superior occipital gyrus | NS-t/FA-w |
| R cuneus - L superior occipital gyrus | NS-t/FA-w |

#### 3.2.5 Reproducibility of graph theoretical metrics

Fig. 9 shows, the ICC distributions over the 90 nodes of the graphs, for the four local graph theoretical metrics: node degree, clustering coefficient, betweenness centrality and local efficiency. The NS-thr scheme resulted in the ICC distributions with the highest mean for all these metrics, with the distributions also appearing the tightest around the mean (the only possible exception being the distribution for the betweenness centrality of the NS OMST scheme). The NS-t/FA-w and NS-t/MD-w schemes also resulted in high ICCs for the node degree and the clustering coefficient.

**Figure 9:**
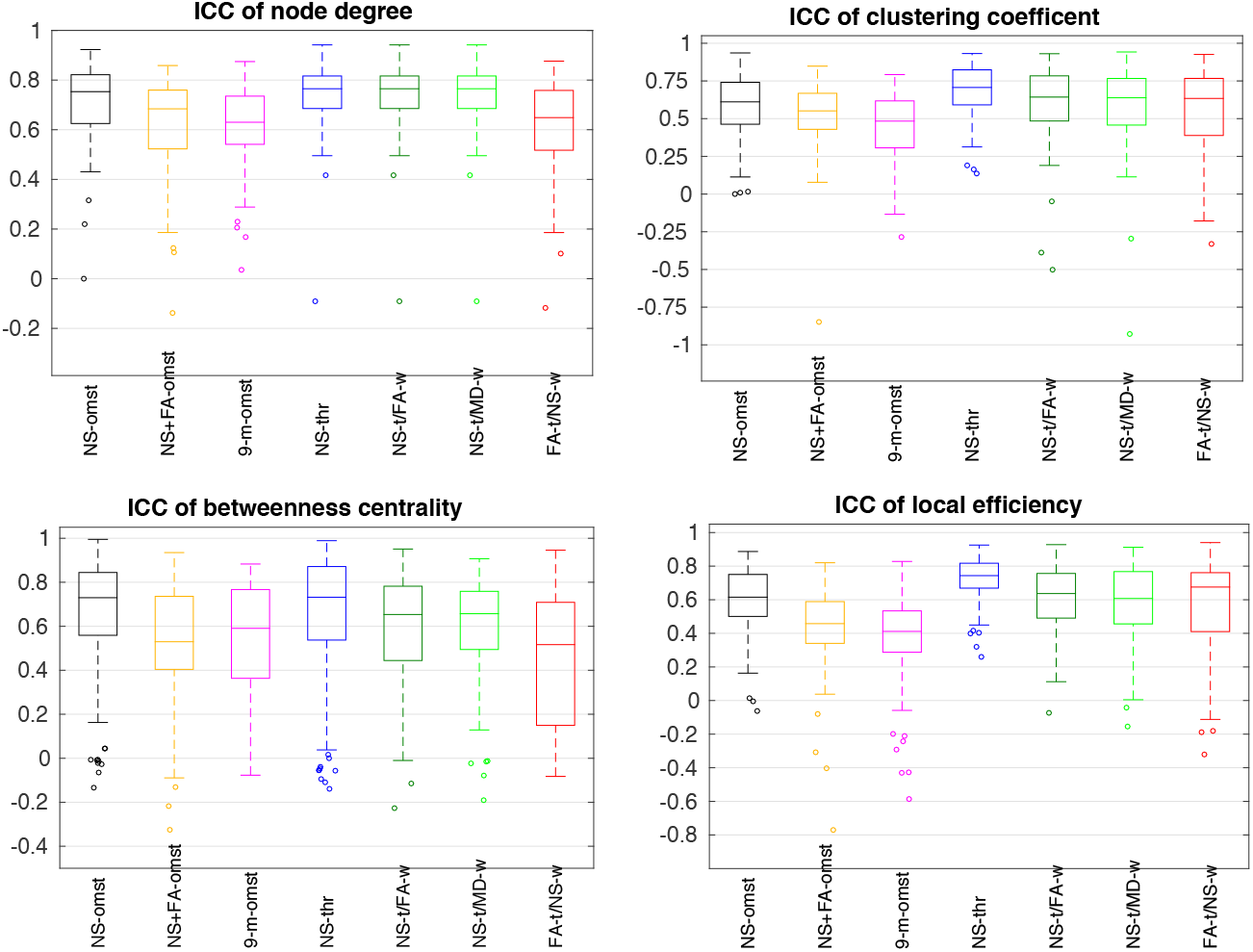
ICCs for the local graph theoretical metrics (*b* = 2000 s/mm^2^). In each box, the middle line shows the mean of the distribution, the edges of the box represent the 25% and 75% points, and the circles indicate points that are thought to be outliers. Each distribution is over the 90 nodes of the graph.

Fig. 10 shows the distributions of the absolute value of the fractional difference between the two scans for the same graph theoretical metrics, for the seven schemes. The 9-m OMST scheme had the lowest mean values and very tight distributions around those means, for all four node-level graph theoretical metrics considered. It is noteworthy that most schemes resulted in comparable distributions for the absolute fractional differences, with the FA-t/NS-w scheme, and the NS OMST scheme for the case of the local efficiency and the clustering coefficient, being the exceptions.

**Figure 10:**
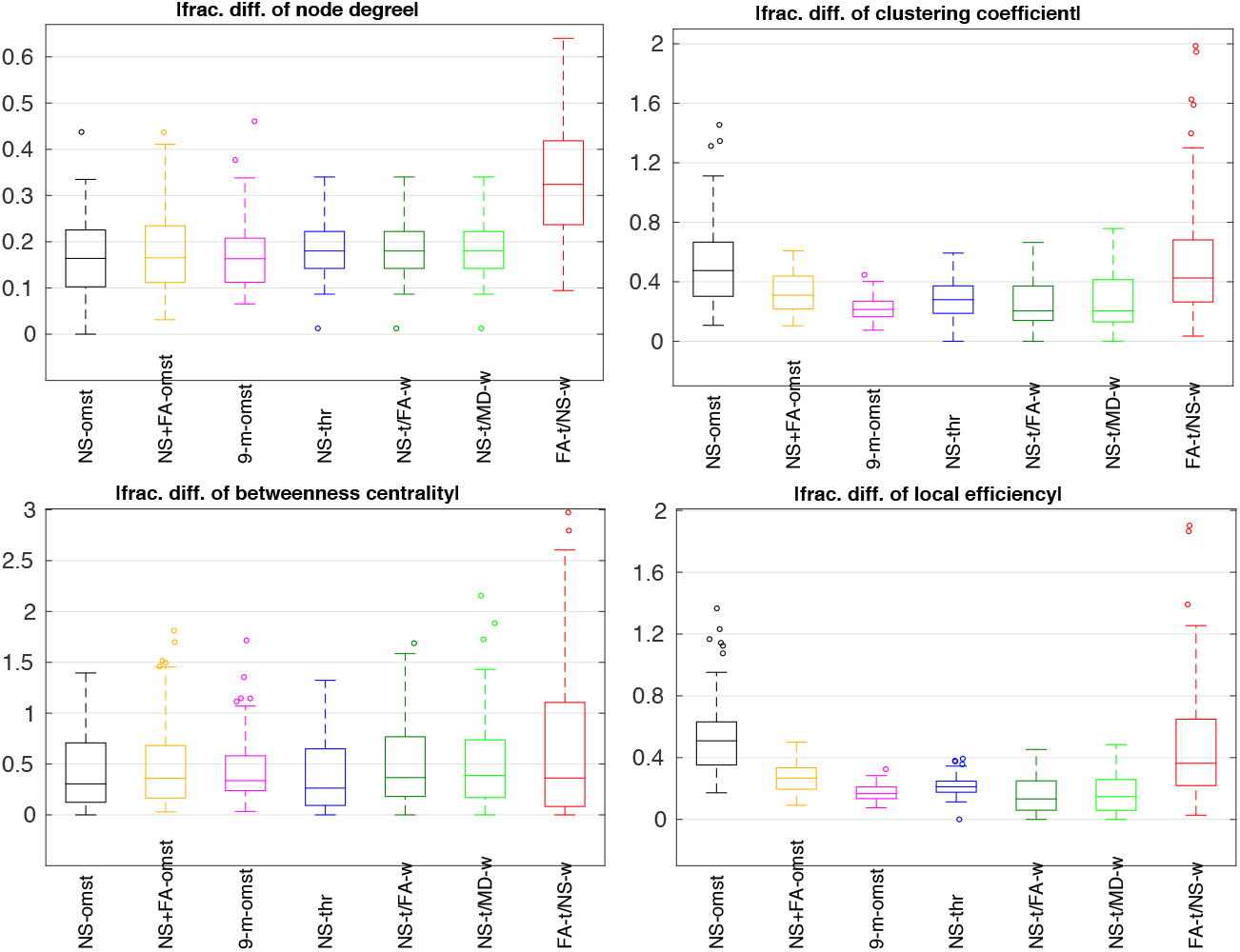
Absolute fractional differences between the two scans, for the local graph theoretical metrics (*b* = 2000 s/mm^2^). In each box, the middle line shows the mean of the distribution, the edges represent the 25% and 75% points, and the circles indicate points that are thought to be outliers. Each distribution is over the 90 nodes of the graph.

The ICCs and mean absolute fractional differences of the global efficiency and the characteristic path length are given in Table 10. Most schemes resulted in low values for the mean absolute fractional differences, however the ICCs for both the global efficiency and the characteristic path length were above 0.85 only for the schemes that rely exclusively on the NS to construct the graphs.

**Table 10:** ICCs and mean absolute fractional differences for the global efficiency and the characteristic path length, for the different schemes, for *b* = 2000 s/mm^2^.

|  |  | NS OMST | NS+FA OMST | 9-m | NS-thr | NS-t/FA-w | NS-t/MD-w | FA-t/NS-w |
| --- | --- | --- | --- | --- | --- | --- | --- | --- |
| $E_{\text{glob}}$ | ICC | 0.86 | 0.23 | 0.64 | 0.87 | 0.58 | 0.64 | 0.50 |
|  | \| frac. diff. \| | 0.076 | 0.031 | 0.059 | 0.077 | 0.063 | 0.064 | 0.173 |
| char path length | ICC | 0.86 | 0.30 | 0.72 | 0.87 | 0.51 | 0.57 | 0.79 |
|  | \| frac. diff. \| | 0.073 | 0.068 | 0.087 | 0.084 | 0.080 | 0.084 | 0.131 |

### 3.3 Power of Structural Network Studies

Consider a study that aims to evaluate whether an intervention performed in a group of participants results in an overall change in their structural networks. Assume that the group is split into two subgroups with equal number of participants *N*, with the intervention performed in only one of the two groups, for example group 2, and no intervention is performed on group 1.

#### 3.3.1 Impact on graph changes

The graphs before and after the intervention are constructed based on a given scheme and the graph similarities are calculated. The distributions of the graph similarities for the two groups will have different means, differing by *µ*, and the same SD *σ*. For the dataset used in this work, the NS-thr scheme resulted in the graph similarity distribution with the lowest SD (Table 4). Table 11 shows the ratio of the SD for the graph similarity distribution of each scheme over the SD of the NS-thr scheme, as well as the factor by which the number of participants needs to be multiplied in order to maintain the same power as when using the NS-thr scheme, based on keeping the quantity Q from Eq. (2.1) constant. The increase in the required number of participants can be very significant if a non-optimal scheme is chosen. For example, if the scheme with the second smallest SD is chosen, namely the NS-t/FA-w scheme, nearly five times as many participants would be required to observe a difference in the graph similarity between the two groups, and therefore to ascertain a possible effect of the intervention, for a given required power and statistical significance.

**Table 11:** Ratio of SD of graph similarity distribution for a given scheme over the SD for the NS-thr scheme. Also given is the increase in the number of participants required to achieve the same power of identifying a difference between populations, assuming normal distributions with different means.

|  | NS OMST | NS+FA OMST | 9-m OMST | NS-t/FA-w | NS-t/MD-w | FA-t/NS-w |
| --- | --- | --- | --- | --- | --- | --- |
| **ratio of SDs** | 2.54 | 3.36 | 2.77 | 2.21 | 2.48 | 5.58 |
| **required increase in number of participants** | 6.5 | 11.3 | 7.7 | 4.9 | 6.2 | 31.2 |

#### 3.3.2 Impact on graph theoretical metrics

The distributions (over the 90 nodes of each graph) of the SDs of the local graph theoretical metrics for data acquired with *b* = 2000 s/mm^2^ are shown in Fig. 11. The distributions of the SDs for the node degree and the betweenness centrality are given in a logarithmic scale. The scheme that exhibits the lowest SD for different graph theoretical metrics depends on the metric itself, and that prevents us from giving a recommendation on the best scheme to use for all these metrics. However, both the 9-m OMST and the NS-thr schemes result in small SDs for these metrics, so would be good choices.

**Figure 11:**
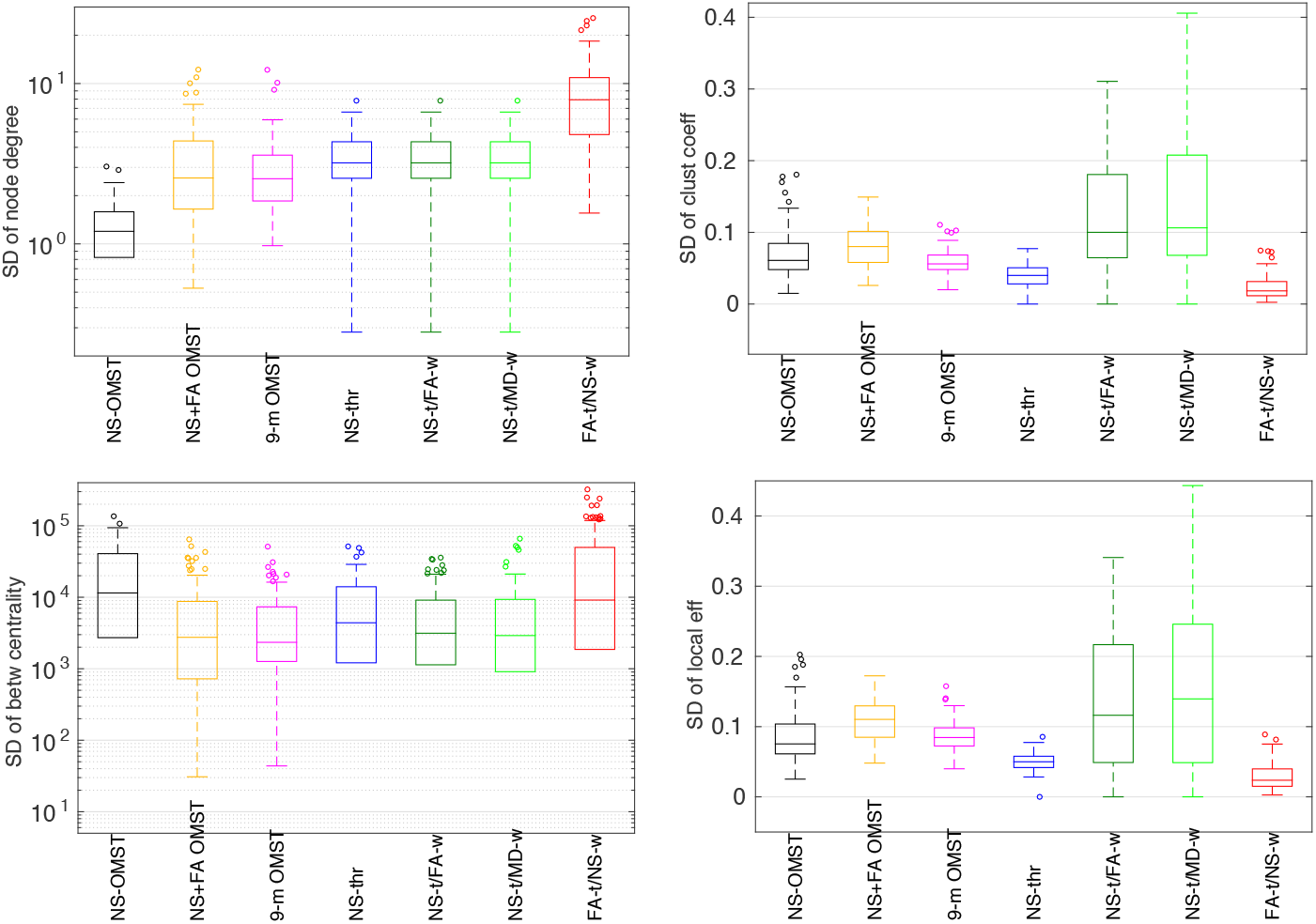
Standard deviation of the local graph theoretical metrics for the seven schemes (first scan, *b* = 2000 s/mm^2^). In each box, the middle line shows the mean of the distribution, the edges of the box represent the 25% and 75% points, and the circles indicate points that are thought to be outliers.

The values of the SDs for the global metrics are given in Table 12. Considering again the study mentioned at the start of this Section, Table 12 implies that if, for example, the NS-thr scheme is used instead of the NS-OMST one, the number of participants needed to detect a change in the characteristic path length, which scales as the square of the ratio of the SDs of the two schemes, would have to be (0.003*/*0.002)^2^ = 2.25 times larger. This can be a very significant difference, specifically for interventions on patient populations, for which recruitment of participants could be challenging.

**Table 12:** SDs for the global efficiency and the characteristic path length for the graphs from each scan, for DWIs acquired with *b* = 2000 s/mm^2^.

|  |  | NS OMST | NS+FA OMST | 9-m | NS-thr | NS-t/FA-w | NS-t/MD-w | FA-t/NS-w |
| --- | --- | --- | --- | --- | --- | --- | --- | --- |
| **E<sub>glob</sub>** | S1 | 0.013 | 0.007 | 0.025 | 0.014 | 0.023 | 0.028 | 0.006 |
|  | S2 | 0.011 | 0.009 | 0.014 | 0.011 | 0.025 | 0.030 | 0.006 |
| **char path length** | S1 | 0.002 | 0.004 | 0.009 | 0.003 | 0.008 | 0.009 | 0.002 |
|  | S2 | 0.002 | 0.004 | 0.006 | 0.003 | 0.008 | 0.009 | 0.002 |

### 3.4 Comparison of Graphs Resulting From Different Diffusion Weightings

As noted earlier, DWIs acquired with different diffusion weightings have different benefits and drawbacks as far as parameter fit, tractography results, etc are concerned. It is important to understand whether the graphs generated through the same scheme using DWIs acquired with different diffusion weightings are similar to each other. Fig. 12 shows the graph similarity for the three pairs of *b*-values compared.

**Figure 12:**
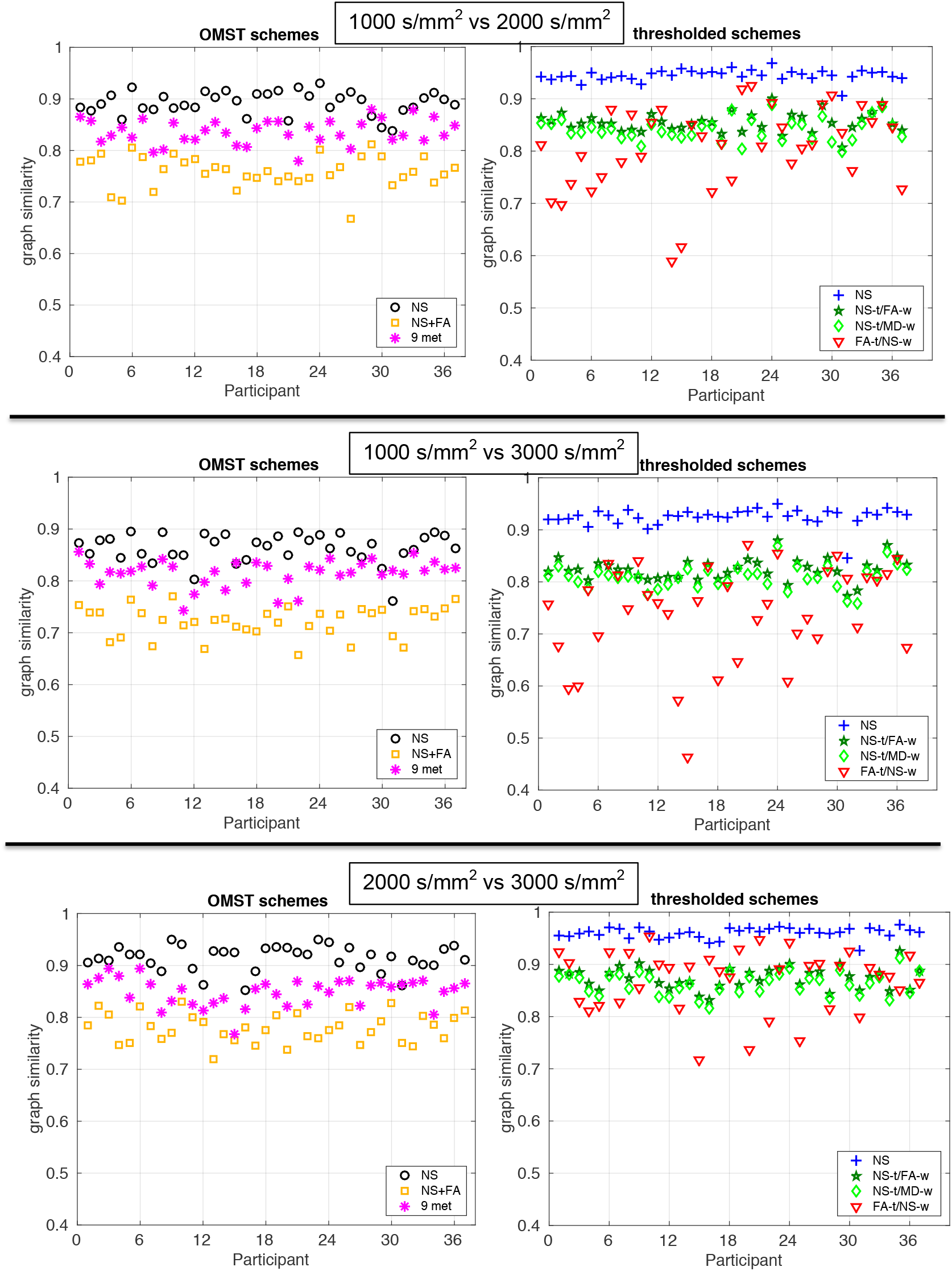
Comparison of graphs across diffusion weightings. The top row shows the plots for comparison between *b* = 1000 s/mm^2^ and *b* = 2000 s/mm^2^, the middle row between *b* = 1000 s/mm^2^ and *b* = 3000 s/mm^2^, and the bottom row *b* = 2000 s/mm^2^ and *b* = 3000 s/mm^2^. Each point gives the similarity of the graphs generated with the same scheme using the data from two different *b* values. For ease of comparison, the vertical axes of all the plots are the same.

In most cases, the graphs generated from DWIs of different *b*-values were very similar to each other, with values of the graph similarity of above 0.8. The FA-t/NS-w scheme exhibited very variable graph similarity between different *b*-values, in particular between *b* = 1000 s/mm^2^ and *b* = 3000 s/mm^2^. This is likely the result of the fact that the FA, which is used to define the topology in that scheme, is not as reliably measured with the DWIs of *b* = 3000 s/mm^2^ as it is with DWIs of smaller *b*. Of the OMST schemes, the NS+FA scheme had the lowest graph similarity between graphs constructed with data acquired at different *b*-values. The highest similarity was exhibited by the NS-thr graphs.

## 4 Discussion

We presented a study on the reproducibility of graphs and their various attributes for different graph-construction schemes, for structural brain networks in the human brain, using the HCP test-retest data. They key findings of our work have implications on longitudinal and comparative studies, as well as on studies that combine dMRI data collected with different diffusion weightings. They are as follows:

1. Different graph-construction schemes result in networks with distinct topologies and edge weights. Even though not surprising, this is an important point because the structural connectome supports function in the brain (Bassett and Gazzaniga, 2011; Mill et al., 2017) and connections between nodes in structural networks can be used to understand the mechanisms that underlie functional connectivity. This is relevant not only for whole-brain studies, but also, and maybe even more so, for studies that use structural sub-networks to study the human brain (for example Drakesmith et al. (2015)).
2. The reproducibility of the graphs depends on the graph construction scheme, with some schemes exhibiting mean similarity of the order of 0.9 or higher while others falling in the range of about 0.8. For the data used in this study, the NS-thr scheme gave the highest mean graph similarity, for all three diffusion weightings. Additionally, the SD of the graph similarity depends on the graph-construction scheme used. Knowing both the mean and the SD of the similarity distribution for healthy participants in the absence of ageing or intervention is essential, because it allows one to make reliable predictions for the number of participants required to observe changes in structural brain networks in developmental, intervention or patient studies.
3. The reproducibility of the topology of graphs depends on the graph construction scheme. The NS-OMST scheme gave the highest topology reproducibility, for all three diffusion weightings. This is in contrast to the fact that the NS-thr scheme resulted in the highest reproducibility for the weighted graphs, and implies that studies which focus on the topology of structural networks rather than on the specific weights of the edges, or studies which use binary rather than weighted networks, may benefit from using different schemes to construct the graphs representing the structural networks.
4. The graphs constructed from DWIs of different diffusion weightings exhibit very good similarity with each other, for most schemes. Of the schemes we investigated in this work, the schemes that rely exclusively on the NS are the most reliable across diffusion weightings. The good similarity implies that each specific combination of algorithms used to fit the DT, perform the tractography and construct the graphs, reveals a fairly consistent picture of the structural connectome, regardless of the diffusion weighting used to acquire the data. The only possible exception is the FA-t/NS-w scheme, which results in graphs that exhibit large deviations from each other, in particular between diffusion weightings of *b* = 1000 s/mm^2^ and *b* = 3000 s/mm^2^, with the graph similarity reaching values of under 0.65 for several participants. This could be due to the limited validity of the DT formalism for *b*-values above 2000 s/mm^2^, in combination with the limited capability of DWIs with *b* = 1000 s/mm^2^ to give good tractography results for crossing fibers. The NS+FA OMST scheme also shows limited reliability when comparing graphs generated from DWIs with *b* = 1000 s/mm^2^ and *b* = 3000 s/mm^2^. This also argues for a possible role that the limited accuracy of the calculation of the FA for larger *b*-values plays in the reproducibility of those graphs across diffusion weightings.
5. The weights of edges that show up in the graphs from both scans of participants were more reliably reproduced for the 9-m OMST, NS-t/FA-w and NS-t/MD-w schemes. The GLM analysis revealed that the absolute difference in edge weights between scans correlates with different WM tract metrics for the different schemes. The NS-t/FA-w and NS-t/MD-w schemes also resulted in the smallest percentage of edges reproduced in the graphs from only one of the two scans of the participants. Such edges are generally characterized by low TV and low NS, although we did not find any statistically significant differences in the distributions of those two quantities between edges that are and are not reproduced in the graphs from both scans. These results give some insight into the attributes of edges that are highly reproducible. More importantly, they highlight the fact that, regardless of the graph construction scheme used, not all edges of structural networks are equally reliable.
6. The reproducibility of graph theoretical metrics depends on the graph construction scheme. The NS-thr, NS-t/FA-w and NS-t/MD-w schemes resulted in the ICC distributions over nodes with the highest means. The 9-m OMST scheme resulted in the lowest mean for the distributions of the absolute fractional differences of the node degree, clustering coefficient and the local efficiency, also resulting in a very tight distribution for the 90 nodes. The results held for the graphs constructed using the DWIs of different diffusion weightings. For the global graph theoretical metrics, the NS-OMST and NS-thr schemes exhibited the highest ICCs, while the NS+FA-OMST scheme exhibited the lowest mean absolute fractional differences.

In addition to these important findings, we showed the effect that a non-optimal choice of scheme can have on the power of studies to detect differences in populations or effects of interventions. Our analysis makes it clear that a careful assessment of the impact of schemes on the SDs of various metrics can guide the choice of scheme. Since each scheme has its own strengths and weaknesses, such an analysis can also allow a considered choice of metrics to look at, possibly excluding metrics that have distributions with large SDs and which are expected to not be sufficiently informative or powerful. In addition to saving time and resources, this can reduce the need for multiple comparison corrections, additionally strengthening the performed analyses.

Furthermore, our results go some way into proving why different studies lead to incongruent conclusions for the network characteristics of patient populations or for the developmental or interventional changes observed in structural brain networks. Even if they use the same acquisition protocols and diffusion weightings for data collection, studies that use different schemes to construct the graphs representing the structural networks are bound to give different results for the topologies and for the significance of the edges. They are also bound to exhibit different reproducibility for the graph theoretical metrics and therefore different capability to observe changes or differences in them. In other words, each scheme is “tuned into” specific graph theoretical metrics and is best suited to detect differences or changes in those.

One of the main goals of brain network analyses is to understand the structural underpinnings of the functional organization of the human brain. Functional networks generated from magnetoencephalography or electroencephalography data have revealed that the strength of functional connections between brain areas depends on the frequency of the signals (Brookes et al., 2011; Deco et al., 2017; Messaritaki et al., 2017; Tewarie et al., 2019). Additionally, they have shown that there may be more than one mechanism of coupling between brain areas, such as phase-phase, phase-frequency, phase-amplitude and amplitude-amplitude (Hyafil et al., 2015; Dimitriadis et al., 2015; Dimitriadis and Salis, 2017; Dimitriadis, 2018; Dimitriadis et al., 2018). Given the distinct mechanisms that the different frequencies and ways of coupling imply, it is possible that different characteristics of the WM tracts support each mechanism. Therefore, different graph construction schemes could result in structural network graphs that have higher overlap with the different functional networks, and can therefore be better suited to understand the corresponding mechanisms (Tewarie et al., 2019). This point is further solidified by our finding that different edges appear to be strongest for each graph-construction scheme.

Our study has a few limitations which we now discuss. The main limitation is that the specific results relating to the optimal scheme to use apply specifically to the HCP dataset we used, and cannot be generalized to other datasets that may involve different scanners or acquisitions, or to analysis strategies that involve different tractography algorithms. The point is clear, however, that it is very much worth looking into the optimal scheme for graph construction for each study, because such an investigation can yield a significant gain in the power of the study. Additionally, the reproducibility was assessed on the basis of two scans. Three or more scanning sessions would give a more robust assessment of the reproducibility. Furthermore, scanning the participants in a controlled manner, making sure that they are scanned at the same time of day, for example, would also be desirable. Lastly, our thresholded networks used a threshold so that their sparsity is the same as the sparsity of the OMST networks. Different thresholds could lead to different results for the reproducibility of the graphs and their attributes for those schemes, and could also be worth exploring.

## 5 Conclusions

We presented a comprehensive analysis of the dependence of the reproducibility of graphs and their various attributes on the graph-construction schemes used. The graph-construction schemes had a significant impact on the reproducibility and, as a result, on the power of the studies to detect differences between populations, maturation or intervention effects. In order for studies to achieve the maximum power in detecting such effects, similar analysis need to always be performed at the outset.

## 6 Acknowledgements

We are grateful to the Human Connectome Project for making the test-retest data freely available. The work was partly funded under the BRAIN Biomedical Research Unit, which is funded by the Welsh Government through Health and Care Research Wales. DKJ is supported by a Wellcome Trust Investigator Award (096646/Z/11/Z) and a Wellcome Trust Strategic Award (104943/Z/14/Z). SID was supported by MRC grant MR/K004360/1 and a MARIE-CURIE CO-FUND EU-UK Research Fellowship.

